# Resetting proteostasis with ISRIB prevents pulmonary fibrosis

**DOI:** 10.1101/2020.02.26.965566

**Authors:** Satoshi Watanabe, Nikolay S. Markov, Ziyan Lu, Raul Piseaux Aillon, Saul Soberanes, Constance E. Runyan, Ziyou Ren, Rogan A. Grant, Mariana Maciel, Hiam Abdala-Valencia, Yuliya Politanska, Kiwon Nam, Lango Sichizya, Hermon G. Kihshen, Nikita Joshi, Alexandra C. McQuattie-Pimentel, Richard I. Morimoto, Paul A. Reyfman, G.R. Scott Budinger, Alexander V. Misharin

**Affiliations:** Department of Medicine, Division of Pulmonary and Critical Care Medicine, Northwestern University Feinberg School of Medicine, Chicago, IL, 60611, USA; Department of Respiratory Medicine, Kanazawa University Graduate School of Medical Sciences, Kanazawa, 920-8641, Japan; Deparment of Molecular Biosciences, Northwestern University, Evanston, IL 60208

**Keywords:** pulmonary fibrosis, aging, integrated stress response, ISRIB

## Abstract

Aging is among the most important risk factors for the development of pulmonary fibrosis. We found that a small molecule that specifically inhibits translational inhibition induced by activation of the integrated stress response (ISRIB) attenuated the severity of pulmonary fibrosis in young and old mice. The more severe fibrosis in old compared to young mice was associated with increased recruitment of pathogenic monocyte-derived alveolar macrophages. Using genetic lineage tracing and transcriptomic profiling we found that ISRIB modulates stress response signaling in alveolar epithelial cells resulting in decreased apoptosis and decreased recruitment of pathogenic monocyte-derived alveolar macrophages. These data support multicellular model of fibrosis involving epithelial cells, pathogenic monocyte-derived alveolar macrophages and fibroblasts. Inhibition of the integrated stress response in the aging lung epithelium ameliorates pulmonary fibrosis by preventing the prolonged recruitment of monocyte-derived alveolar macrophages.

## INTRODUCTION

Pulmonary fibrosis is a relentlessly progressive and often fatal disease characterized by the replacement of normal lung tissue with fibrotic scar that increases the work of breathing and impairs gas exchange (1, 2). Advanced age is among the most important risk factors for the development of pulmonary fibrosis (3). Even in patients with a genetic predisposition, the onset of disease seldom occurs before the sixth decade and the incidence increases exponentially with advancing age (4, 5). The development of cellular senescence, mitochondrial dysfunction, stem cell exhaustion and proteostatic stress have all been implicated in the age-related susceptibility to pulmonary fibrosis, but the molecular mechanisms linking these pathways with the development of fibrosis are poorly understood (6, 7).

Proteostasis refers to the dynamic process by which cells control the concentration, conformation, binding interactions, and stability of individual proteins making up the proteome via regulated networks that influence protein synthesis, folding, trafficking, disaggregation, and degradation (8, 9). Analyses of families with high rates of pulmonary fibrosis as well as unbiased genetic studies in populations of patients with idiopathic pulmonary fibrosis (IPF) identified associations between mutations in genes predicted to disrupt proteostasis in the alveolar epithelium and the development of pulmonary fibrosis (10). Proteostatic stress activates the intracellular sensors PERK, PKR, GCN2 or HRI to induce the integrated stress response (ISR) (11). Activation of the ISR reduces the rate of global protein translation while enhancing the translation of selected transcription factors, including ATF4 and CHOP, that induce the expression of chaperones and other cell-protective genes. Prolonged or high-level induction of CHOP activates apoptotic pathways that result in cell death (12). The ISR is also activated via PKR in response to viral infections, which have been implicated in exacerbations of IPF (13).

We and others have suggested that proteostatic stress induced by genetic mutations or environmental agents that drive fibrosis leads to the recruitment of profibrotic, monocyte-derived alveolar macrophages to the lung (14–25). We have shown that these monocyte-derived alveolar macrophages are maintained by autocrine or paracrine signaling via M-CSF/M-CSFR and localized specifically to regions of epithelial injury in close proximity to fibroblasts, inducing their activation, proliferation and secretion of matrix proteins (18). Over time, these regulatory circuits between macrophages and fibroblasts may become dispensable, creating autonomous regions of progressive fibrosis driven exclusively by fibroblasts (26, 27). Even so, therapeutic strategies that reduce the ongoing recruitment of monocyte-derived alveolar macrophages and prevent the initiation of macrophage-fibroblast circuits are likely to be effective as the progression of the human disease is heterogenous in space and time.

We found that enhanced and prolonged fibrosis in old compared with young mice was associated with prolonged recruitment of pathogenic monocyte-derived alveolar macrophages suggesting a failure to repair epithelial injury in older animals. A small molecule integrated stress response inhibitor (ISRIB) specifically prevents translational inhibition and therefore other downstream consequences of activation of the ISR and has shown promise as a therapeutic strategy against inflammation, cancer and neurocognitive diseases in aging (28, 29). We found that ISRIB attenuated the development of fibrosis in young and aged animals in resolving (bleomycin) and non-resolving (asbestos) models of pulmonary fibrosis. The administration of ISRIB ameliorated pulmonary fibrosis by increasing the resilience of epithelial cells to proteostatic stress, preventing their apoptosis and attenuating the recruitment of pathogenic monocyte-derived alveolar macrophages.

## RESULTS

### Aging is associated with more severe pulmonary fibrosis in response to bleomycin and asbestos

We induced pulmonary fibrosis in young mice (3-months) and old mice (over 18-months) via the intratracheal administration of bleomycin (0.025 unit/50 µl), and analyzed the lungs 28 days later (Figure 1A-D). Old mice developed more severe pulmonary fibrosis compared with young mice, as evidenced by decreased lung compliance, elevated collagen levels, and worsened lung pathology (Figure 1A and B). While collagen levels in young mice returned to the near-normal levels, increased collagen levels persisted in old mice for up to 56 days (Figure 1C). Both young and old mice lost body weight after the instillation of bleomycin. However, while body weight recovered in young mice, it failed to recover in old animals (Figure 1D).

**Figure 1.**
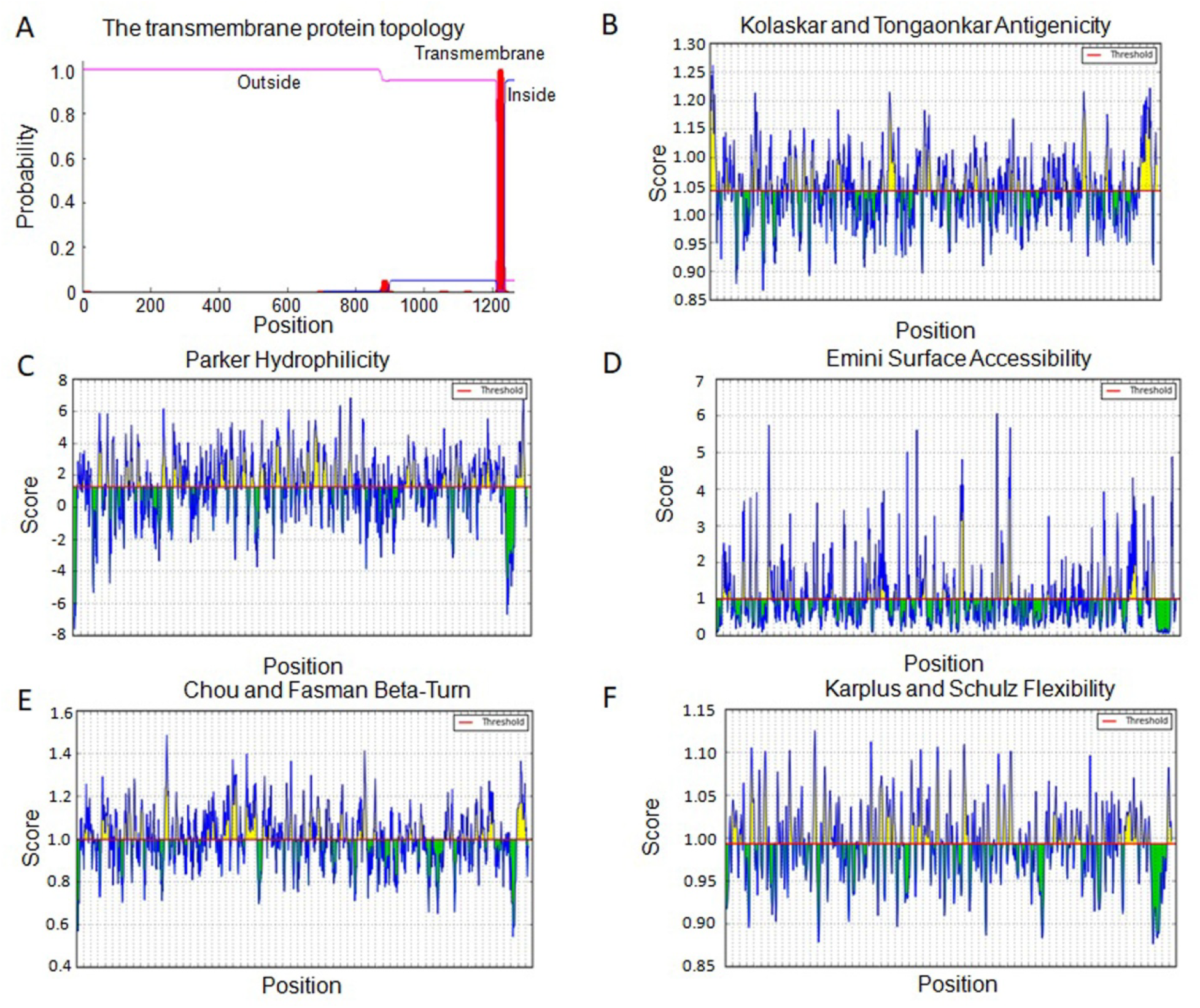
Old mice develop more severe lung fibrosis compared to young mice. Young mice (3 months) and old mice (over 18 months) were administered bleomycin (0.025 unit/50 µl) or crocidolite asbestos (100 μg/50 µl) intratracheally and the lungs were harvested 28 days later. (**A**) Representative histologic images in bleomycin model (Masson’s trichrome, Scale bar = 1 mm). (**B**) Quantification of fibrosis score using Ashcroft score, lung compliance using Flexivent, and soluble collagen in lung homogenates in bleomycin model. (**C**) Quantification of soluble collagen in naïve mice and 14, 28, and 56 days after the administration of bleomycin. (**D)** Body weight in bleomycin model with young and old mice. (**E**) Representative histologic images in asbestos model (Masson’s trichrome, Scale bar = 1 mm). (**F**) Quantification of fibrosis score, lung compliance using Flexivent, and soluble collagen in lung homogenates in asbestos model. (**G**) Quantification of soluble collagen in naïve mice and 28 and 56 days after the administration of asbestos. (**H**) Body weight in asbestos model with young and old mice. Data presented as mean ± SEM, 5–10 mice per group, one-way ANOVA with Tukey-Kramer test for multiple comparisons or Kruskal-Wallis test with Dunn’s multiple comparisons test for comparison of each body weight to baseline (day 0). *, p < 0.05. Representative data from two independent experiments.

We observed a similar age-related susceptibility in a non-resolving model of pulmonary fibrosis induced by the intratracheal administration of crocidolite asbestos (100 µg/50 µl) (18, 30). In young mice exposed to intratracheal asbestos, we observed mild peribronchial fibrosis as evidenced by histological scoring of Masson’s trichrome stained lung sections, but body weight and lung compliance were not changed (Figure 1E-H). In contrast, old animals lost and did not recover body weight and had reduced lung compliance, increased lung collagen levels and increased fibrosis area as evident by Masson’s trichrome staining 28 days after asbestos administration (Figure 1E-H). Collagen levels remained increased up to 56 days after asbestos administration (Figure 1G). Thus, aging is associated with increased severity of fibrosis in both bleomycin- and asbestos-induced models of pulmonary fibrosis.

### Recruitment of pathogenic monocyte-derived alveolar macrophages is increased during pulmonary fibrosis in old mice

We and others have performed causal genetic studies in mice to show that monocyte-derived alveolar macrophages are necessary for the development of bleomycin- and asbestos-induced pulmonary fibrosis independent of circulating monocytes, tissue-resident interstitial macrophages or tissue-resident alveolar macrophages (14–18). Therefore, we asked whether the increased severity of pulmonary fibrosis in old mice is associated with an increased recruitment of monocyte-derived alveolar macrophages. We quantified myeloid cell populations in the lung via flow cytometry 28 days after the administration of bleomycin or asbestos, as previously described (16) (Figure 2 and Figure S1). We used differential expression of Siglec F to separate monocyte-derived alveolar macrophages (CD64^+^Siglec F^low^) from tissue-resident alveolar macrophages (CD64^+^Siglec F^high^) (16, 31). We found that numbers of classical monocytes and lymphocytes were increased in the lungs of old mice both in the steady state and during asbestos-induced fibrosis, as has been reported (Figure 2 and Figure S2) (32). After the intratracheal instillation of either bleomycin or asbestos, the number of interstitial macrophages (CD64^+^Siglec F^−^) and monocyte-derived alveolar macrophages was increased in old compared to young mice (Figure 2). In our gating strategy interstitial macrophages include a mixed population of tissue-resident perivascular and peribronchial macrophages and differentiating monocyte-derived macrophages. Thus, the increased severity of pulmonary fibrosis in old mice was associated with increased numbers of profibrotic monocyte-derived alveolar macrophages.

**Figure 2.**
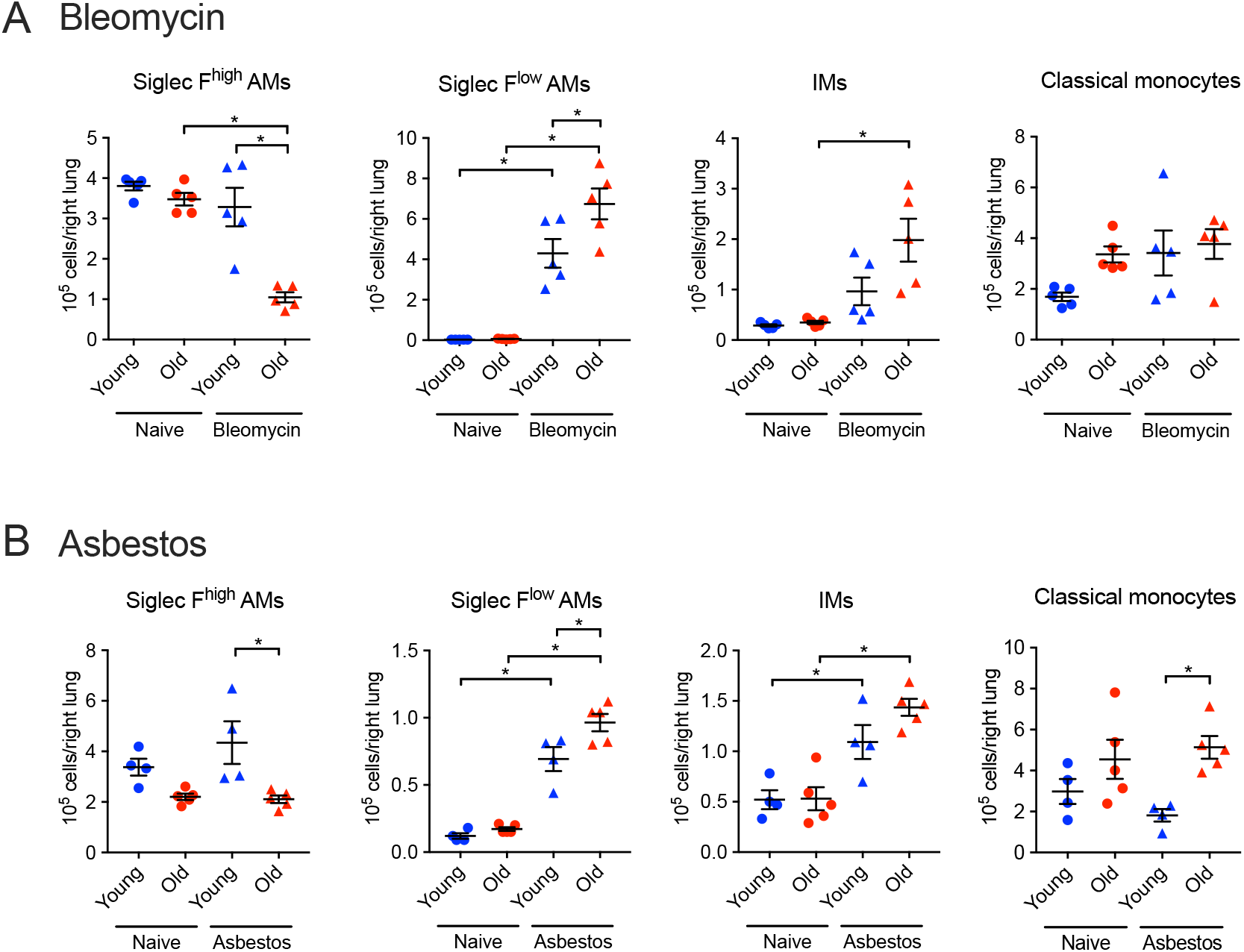
Aging is associated with the increased number of monocyte-derived alveolar macrophages in lung fibrosis. Young mice (3 months) and old mice (over 18 months) were administered bleomycin (0.025 unit/50 µl, intratracheally) or crocidolite asbestos (100 μg/50 µl, intratracheally), and the lungs were harvested 21 days later. Flow cytometry to quantify monocyte and macrophage populations in (**A**) bleomycin model and (**B**) asbestos model. Data presented as mean ± SEM, 5–8 mice per group, one-way ANOVA with Tukey-Kramer test for multiple comparisons. *, p< 0.05. Representative data from two independent experiments.

### ISRIB ameliorates pulmonary fibrosis in young and old mice

Chronic activation of the integrated stress response (ISR) during aging has been linked to the development of pulmonary fibrosis (33–36). We treated young mice with a small molecule inhibitor of the integrated stress response, ISRIB (25 mg/kg via intraperitoneal injections) or vehicle daily starting at day 7 after instillation of bleomycin, when acute lung injury is largely resolved and active fibrogenesis has begun, and continued until harvest at day 28 (Figure 3A). Treatment with ISRIB resulted in higher lung compliance, lower collagen levels and reduced histologic evidence of fibrosis as measured by blinded scoring of randomly selected lung sections stained with Masson’s trichrome (Figure 3C and D). Moreover, young mice treated with ISRIB recovered body weight faster than vehicle-treated mice (Figure 3B). Old mice treated with ISRIB demonstrated attenuated weight loss, increased lung compliance, decreased collagen levels and reduced histologic evidence of fibrosis when compared with vehicle-treated mice (Figure 3E-G). The administration of ISRIB in naïve old or young mice did not affect body weight, lung compliance, collagen levels or induce detectable histologic changes.

**Figure 3.**
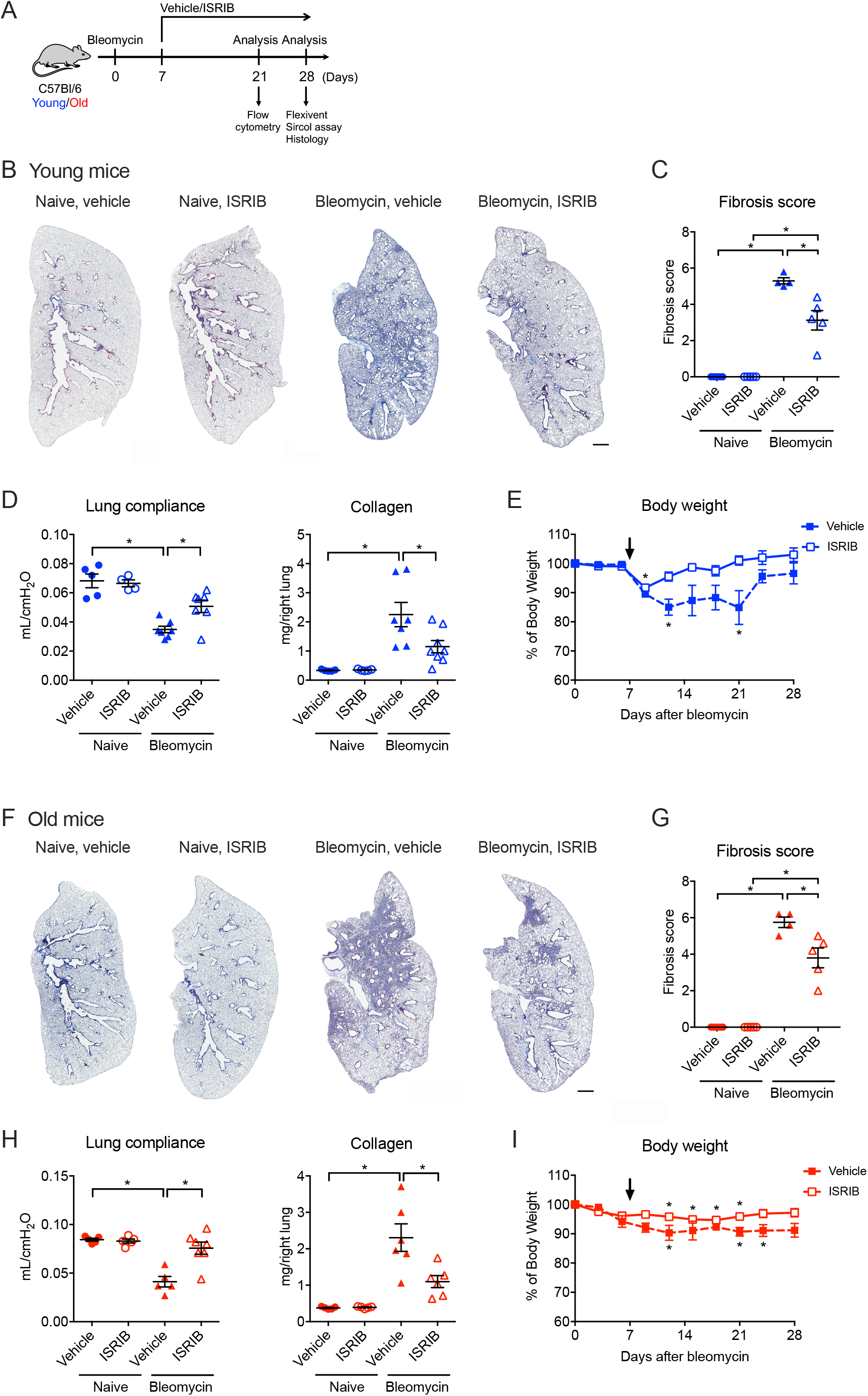
Therapy with ISRIB attenuates bleomycin-induced pulmonary fibrosis in young and old mice. (**A**) Schematic of the experimental design. Young (3 months) and old (over 18 months) mice were administered bleomycin (0.025 unit/50 µl, intratracheally) and treated with 25 mg/kg of ISRIB or vehicle (i.p., daily) beginning at Day 7 when lung injury has resolved and harvested at the indicated time points. (**B**) Representative images of lung tissue from young naïve mice or bleomycin-treated mice with or without ISRIB on day 28. Masson’s trichrome staining. Scale bar = 1 mm. (**C-D**) Quantification of fibrosis score, lung compliance, and collagen levels from young naïve mice or bleomycin-treated mice with or without ISRIB. (**E**) The graph shows the body weight of the young mice. 5 mice per each group. Arrow indicates the starting point of ISRIB treatment. (**F**) Representative images of lung tissue from old naïve mice or bleomycin-treated mice with or without ISRIB on day 28. (**G-H**) Quantification of fibrosis score, lung compliance, and collagen levels from old naïve mice or bleomycin-treated mice with or without ISRIB. (**I**) The graph shows the body weight of the old mice. 5 mice per each group. Arrow indicates the starting point of ISRIB treatment. Masson’s trichrome staining. Scale bar = 1 mm. Data are shown as mean ± SEM, 5-7 mice per group. One-way ANOVA with Tukey-Kramer test for multiple comparison or Kruskal-Wallis test with Dunn’s multiple comparisons test for comparison of each body weight to baseline (day 0). *p < 0.05. Representative data from two independent experiments.

We next tested the therapeutic efficacy of ISRIB in a model of non-resolving fibrosis induced by the intratracheal administration of asbestos. Mice were treated with crocidolite asbestos (100 µg/50 µl), and then treated with ISRIB (25 mg/kg via intraperitoneal injections) or vehicle daily beginning 7 days post-instillation. The lungs were harvested and analyzed 28 days after asbestos treatment (Figure S3A). In both young and old mice treatment with ISRIB was associated with reduced evidence of fibrosis as measured by blinded scoring of Masson’s trichrome stained lung sections (Figure S3B-C and E-F). As the dynamic range of collagen levels using picrosirius red precipitation (Sircol assay) is too small to detect the differences between the groups, we utilized second harmonic generation microscopy for quantitative analysis of collagen (37). Both young and old mice treated with ISRIB showed a decrease in lung collagen compared to vehicle-treated mice (Figure S3D and G). In addition, young mice treated with ISRIB recovered body weight faster than vehicle-treated mice (Figure S3C and F). Together, these data suggest that ISRIB ameliorates pulmonary fibrosis in both young and old mice in both resolving and non-resolving models.

### ISRIB reduces the recruitment of profibrotic monocyte-derived alveolar macrophages to the lung

To test whether the amelioration of fibrosis with ISRIB is associated with decreased recruitment of profibrotic monocyte-derived alveolar macrophages, we treated mice with bleomycin and 7 days later started treatment with ISRIB (25 mg/kg, daily via intraperitoneal injections) followed by analysis of myeloid and lymphoid cells in the lung via flow cytometry at day 21. Interestingly, treatment with ISRIB decreased the number of lung monocytes in old mice, but did not affect monocyte levels in young mice (Figure 4A,B). The administration of ISRIB also decreased the number of interstitial macrophages (CD64^+^Siglec F^−^) and monocyte-derived alveolar macrophages (CD64^+^Siglec F^low^), but did not change the number of tissue-resident alveolar macrophages (CD64^+^Siglec F^high^) in either young or old mice (Figure 4A, B).

**Figure 4.**
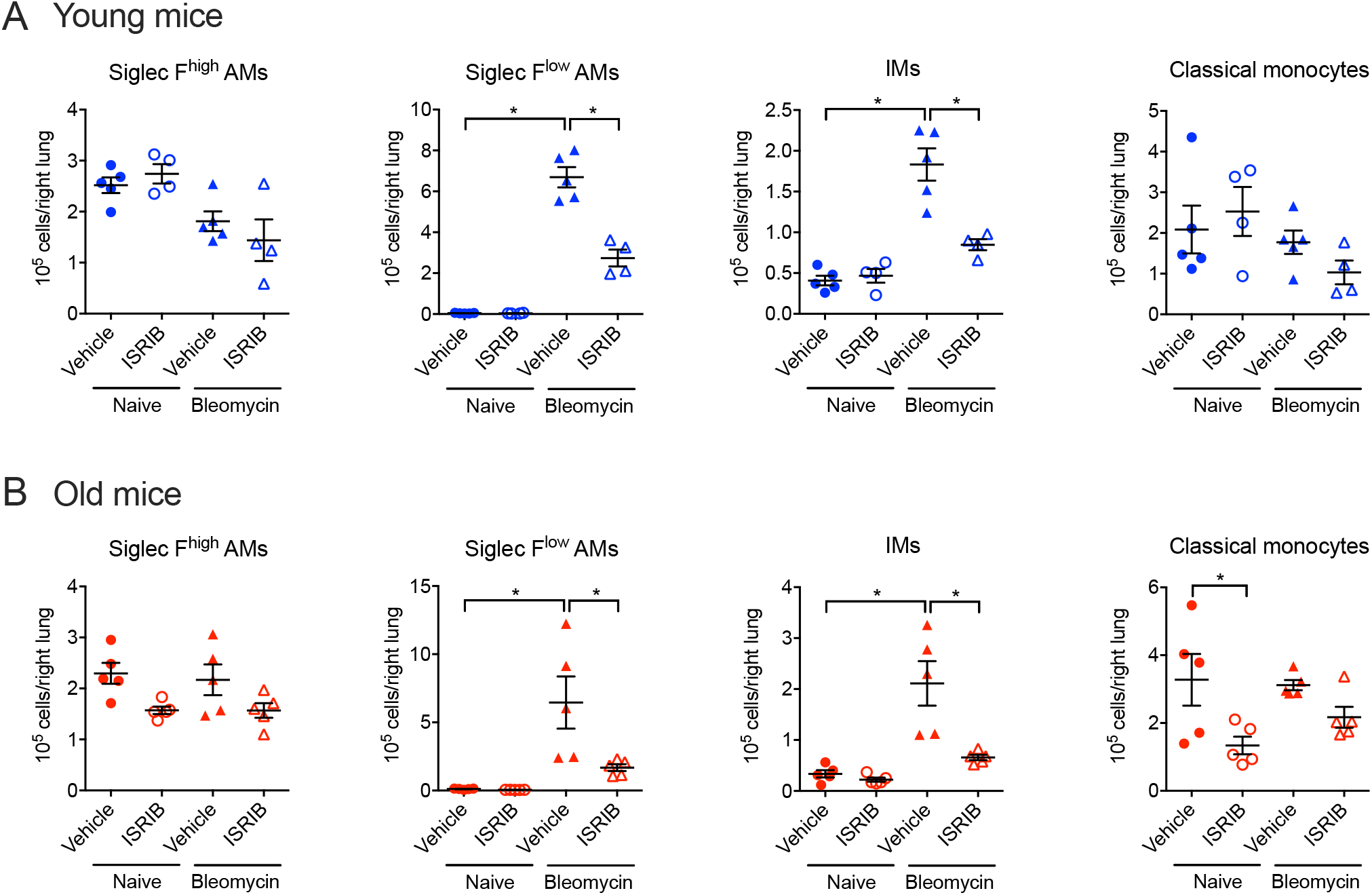
ISRIB reduces the number of monocyte-derived alveolar macrophages in lung fibrosis. Young (3 months) and old (over 18 months) mice were administered 0.025 unit of bleomycin, treated with 25 mg/kg of ISRIB or vehicle (i.p., every day), and the lungs are analyzed by flow cytometry on day 21. The number of macrophages and classical monocytes in (**A**) young and (**B**) old mice. Data are shown as mean ± SEM, 5 mice per group. One-way ANOVA with Tukey-Kramer test for multiple comparison. *p < 0.05. Representative data from two independent experiments.

We used a genetic lineage tracing system to permanently label circulating monocytes and monocyte-derived alveolar macrophages derived from circulating monocytes (*Cx3cr1*^*ERCre*^ × *zsGreen*^*LSL*^). Classical monocytes, which are precursors for monocyte-derived alveolar macrophages, have a half-life of ~5-7 days in mice (38). Therefore, only monocyte-derived alveolar macrophages recruited within this window of time after tamoxifen administration will be labeled with GFP (Figure 5A). We administered tamoxifen simultaneously with bleomycin and started treatment with ISRIB at day 7, harvesting lungs for analysis on days 7, 14 and 21 (Figure 5B). In young mice we found that at all time points nearly 80% of the Siglec F^low^ alveolar macrophages were GFP-positive, suggesting that most of the monocyte-derived alveolar macrophages enter the lung after bleomycin injury as a single wave within the first 7 days with negligible recruitment afterwards (Figure 5C and D). In young mice, the number of Siglec F^low^GFP^+^ alveolar macrophages peaked at day 14 and decreased by day 21. Surprisingly, old mice had fewer Siglec F^low^GFP^+^ alveolar macrophages at day 7 and 14, but more at day 21. The number of Siglec F^low^GFP^−^ alveolar macrophages also increased by day 21, suggesting that persistent epithelial injury in old mice drives the proliferation and ongoing recruitment of monocyte-derived alveolar macrophages (Figure 5E). Treatment with ISRIB did not affect the numbers of tissue-resident alveolar or interstitial macrophages. In contrast, treatment with ISRIB decreased the number of Siglec F^low^GFP^+^ alveolar macrophages at day 14 in young mice and at day 21 in old mice (Figure 5F and G). We have shown that cell surface expression of Siglec F increases as monocyte-derived alveolar macrophages mature (16). Consistently, the expression of Siglec F increased in GFP-positive monocyte-derived alveolar macrophages between days 7 and 21 of bleomycin-induced lung fibrosis in young mice. Interestingly Siglec F levels were higher in GFP-positive macrophages in old mice treated with ISRIB compared with vehicle (Figure S4A and B), suggesting that ISRIB not only reduced recruitment of monocyte-derived alveolar macrophages in old mice but also promoted their maturation.

**Figure 5.**
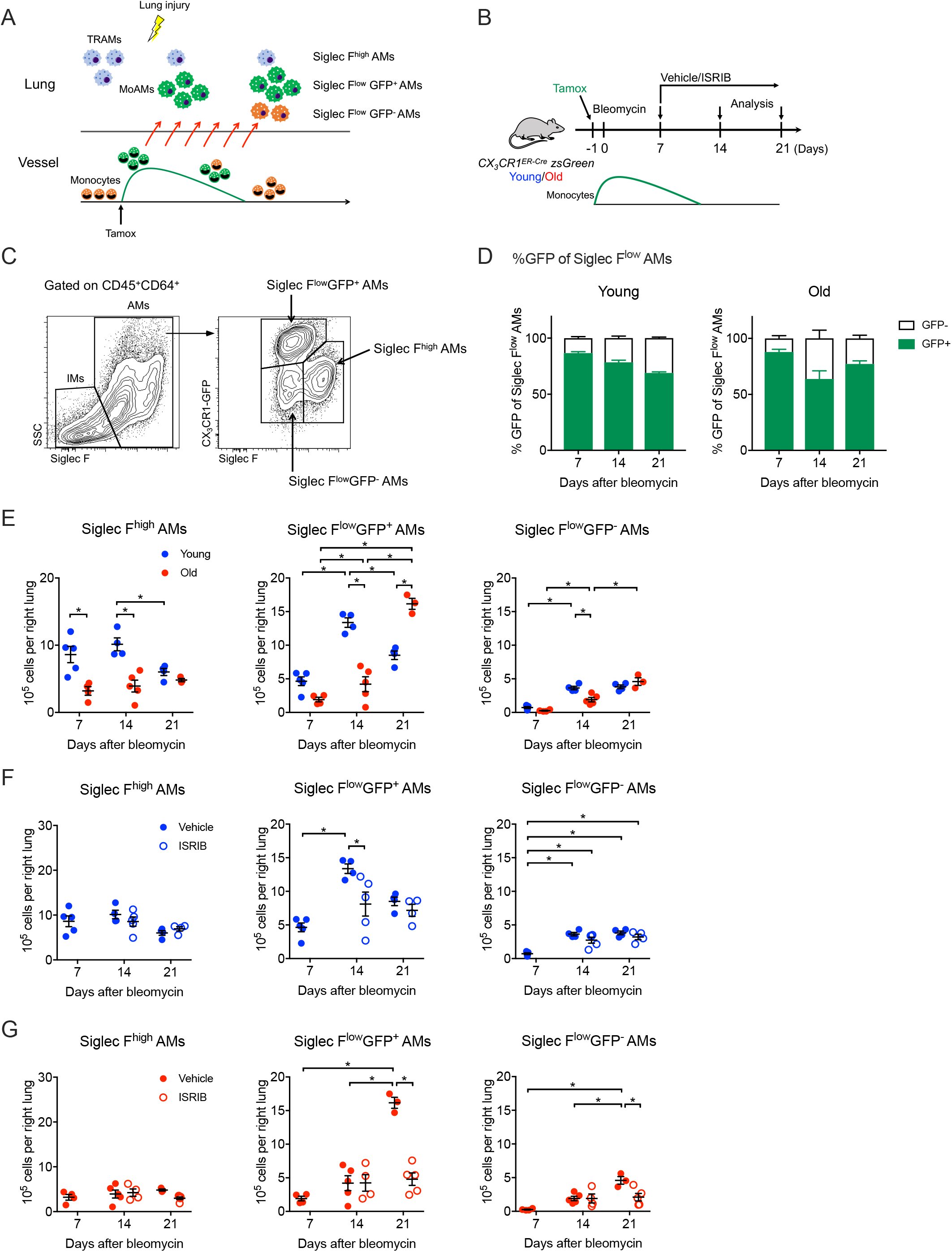
The recruitment kinetics of monocyte-derived alveolar macrophages is different between young and old mice during pulmonary fibrosis. (**A**) Schematic of the genetic lineage tracing system using *CX*_*3*_*CR1*^*ERCre*^ × *zsGreen* mice. Circulating monocytes are labeled with GFP after tamoxifen treatment, but tissue-resident alveolar macrophages are not. After lung injury, GFP-labeled monocytes are recruited to the lung and become monocyte-derived alveolar macrophages. GFP-labeled monocytes are partially replaced by newly formed unlabeled cells 7 days later and are completely replaced 21 days later. In this system, we can track the monocyte-derived alveolar macrophages recruited during the initial wave of monocyte influx. (**B**) Schematic of the experiment design. Young (3 months) and old (over 18 months) female *CX*_*3*_*CR1*^*ERCre*^ × *zsGreen* mice were used to assess the recruited monocyte-derived alveolar macrophages. The mice were given pulse of tamoxifen (10 mg, p.o.) one day before receiving intratracheal bleomycin. Therapeutic ISRIB or vehicle were administered daily beginning at Day 7 after bleomycin and the lungs were harvested at the indicated times. (**C**) Flow cytometry gating strategy for macrophage populations. (**D**) %GFP of Siglec F^low^ alveolar macrophages (AMs) during the course of bleomycin model with young and old mice. (**E**) The number of Siglec F^high^ AMs, Siglec F^low^GFP^+^ AMs and Siglec F^low^GFP^−^ AMs in comparison of young with old mice. (**F-G**) The number of Siglec F^high^ AMs, Siglec F^low^GFP^+^ AMs and Siglec F^low^GFP^−^ AMs in young and old mice in comparison of vehicle with ISRIB treatment. Data are shown as mean ± SEM, 3-5 mice per group. One-way ANOVA with Tukey-Kramer test for multiple comparison. *p < 0.05.

### Transcriptomic profiling suggests that ISRIB promotes the differentiation of monocyte-derived alveolar macrophages

ISRIB reduced number of lung monocytes in old mice, leaving open the possibility that it can directly target monocytes to reduce the recruitment of monocyte-derived alveolar macrophages. We therefore performed RNA-seq on classical monocytes flow-sorted from the lungs of the young and old naïve mice treated with ISRIB for 7 days. Principal component analysis demonstrated a lack of variance in the data related to either aging or treatment with ISRIB (Figure S5A). A direct pairwise comparison between the groups identified only 2 differentially expressed genes between ISRIB- and vehicle-treated mice irrespective of age (Supplemental Table 1). These results suggest that while treatment with ISRIB alters the number of classical monocytes in the lung, it does not substantially modify their transcriptomic identity.

We then performed RNA-seq on flow-sorted monocyte-derived alveolar macrophages in young and old mice with or without treatment with ISRIB (starting at day 7) in both the bleomycin (harvest day 21, Figure 6A, S5B) and asbestos models of lung fibrosis (harvest day 28, Figure 6B, S5C). Principal component analysis demonstrated that treatment with ISRIB and age were the main sources of variance in the dataset (PC1 29.7% and PC2 12.2%, respectively) (Figure S4B). We performed k-means clustering on 1923 differentially expressed genes in the bleomycin dataset (ANOVA-like test, FDR q value < 0.05) (Figure 6A, Supplemental Tables S2 and S3). Treatment of bleomycin exposed young and old mice with ISRIB upregulated genes involved in biosynthetic process, positive regulation of proliferation, and phagocytosis (Figure 6A, Cluster 1) and downregulated genes involved in cell cycle, antigen processing, MHC II presentation and innate immune response (Figure 6A, Cluster 2). Genes that were upregulated in monocyte-derived alveolar macrophages from bleomycin-exposed old compared with young mice included those involved in inflammatory response, antigen processing and presentation, and reactive oxygen species production (Figure 6A, Cluster 4). Genes that were specifically upregulated in monocyte-derived alveolar macrophages from old compared with young mice treated with bleomycin followed by ISRIB (Figure 6A, Cluster 5) were enriched for processes characteristic for homeostatic tissue-resident alveolar macrophages such as lipid metabolism and phagocytosis, while downregulated genes (Figure 6A, Cluster 3) were involved in extracellular matrix remodeling, myoblast proliferation, macrophage migration, and chemotaxis.

**Figure 6.**
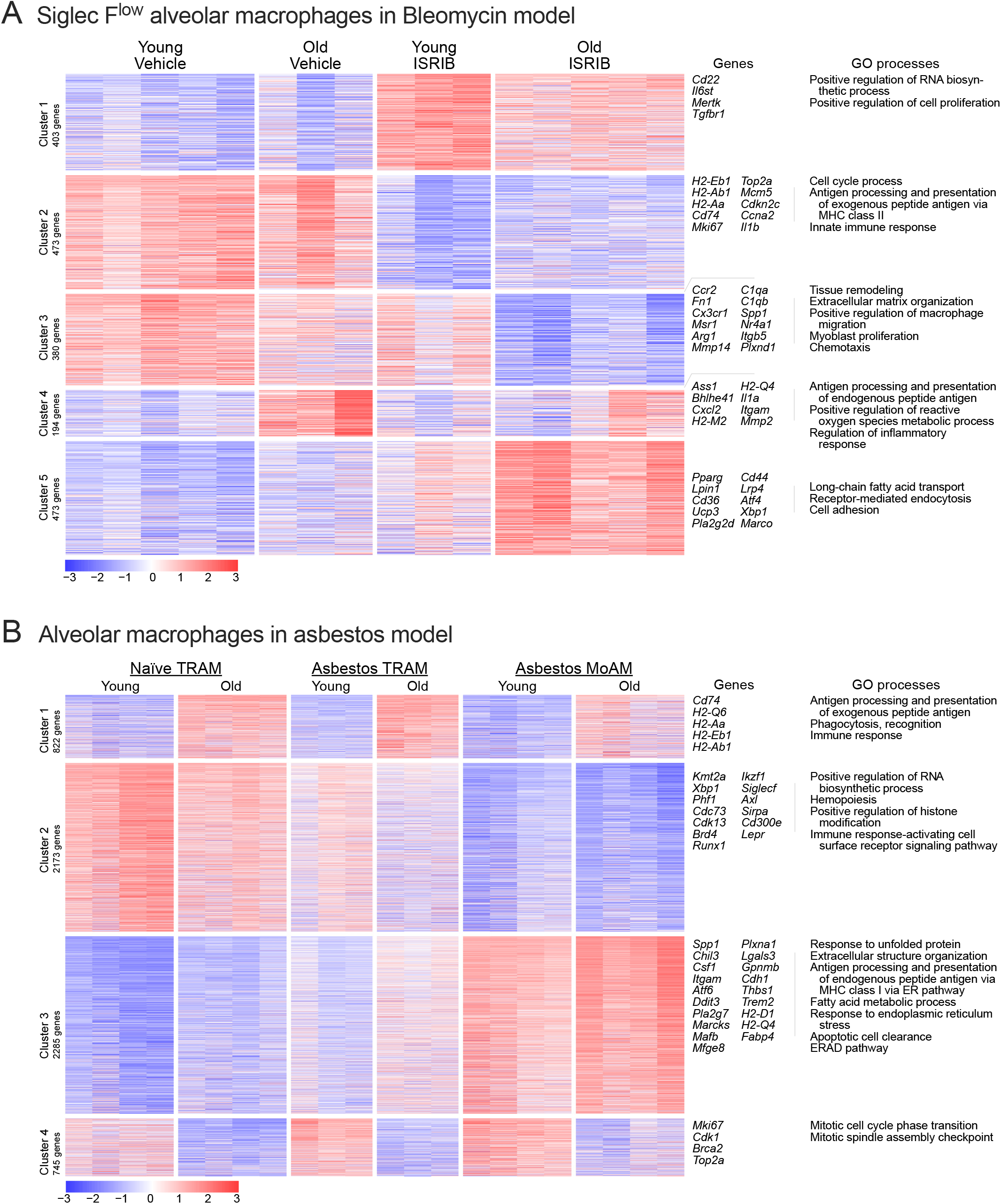
Transcriptomic profiling suggests that ISRIB promotes the differentiation of monocyte-derived alveolar macrophages. (**A**) Transcriptomes of sorted monocyte-derived alveolar macrophages in young and old mice bleomycin model with and without ISRIB treatment. k-means clustering of all identified differentially-expressed genes (ANOVA-like test on negative binomial generalized log-linear model, FDR q-value < 0.05) revealed clusters of genes especially up- and down-regulated by ISRIB treatment (cluster 1 and 2), genes up- and down-regulated by ISRIB especially in old mice (cluster 3 and 5), and genes especially up in old mice and down-regulated by ISRIB (cluster 4). The characteristic GO processes and genes associated with each cluster are shown on the right sides, correspondingly. (**B**) Transcriptomes of sorted tissue-resident alveolar macrophages (TRAMs) and monocyte-derived alveolar macrophages (MoAMs) in naïve lung and asbestos-induced lung fibrosis. k-means clustering of all identified differentially-expressed genes (ANOVA-like test on negative binomial generalized log-linear model, FDR q-value < 0.05) revealed clusters of genes associated with gene ontogeny (clusters 1 and 4) and genes especially up- and down-regulated in MoAMs in asbestos-induced lung fibrosis (clusters 2 and 3). The characteristic GO processes and genes associated with each cluster are shown on the right sides, correspondingly.

Our fate-mapping data suggested that after bleomycin, monocyte-derived alveolar macrophages continued to be recruited much longer after bleomycin in old compared with young mice and this prolonged recruitment was prevented by treatment with ISRIB (Figure 5C,D). These differences in the kinetics of recruitment might account for the apparent reduction in fibrotic gene expression in monocyte derived alveolar macrophages from ISRIB treated animals in the bleomycin model (Figure 6A, Cluster 3). We therefore performed transcriptomic profiling of tissue-resident and monocyte-derived alveolar macrophages in asbestos-induced pulmonary fibrosis, where we have shown that monocytes are continuously recruited (Figure 6B) (18). Principal component analysis demonstrated macrophage ontogeny and aging as two major sources of variance in the dataset (PC1 54.5% and PC2 18.6%, correspondingly, Figure S5C). Importantly, k-means clustering performed on differentially expressed genes (FDR q>0.05 in ANOVA-like test) failed to identify genes that differed between asbestos induced monocyte-derived alveolar macrophages in young compared with old mice (Figure 6B, Supplemental Tables S4 and S5). Instead, genes upregulated in monocyte-derived alveolar macrophages compared with tissue resident alveolar macrophages were similar in young and old animals (Figure 6B, Cluster 3) and included genes related to phagocytosis (*Mefg8, C1qa, C1qb, C3ar, Itgam, Msr1*), extracellular matrix remodeling and fibroblast proliferation (*Itgav, Mmp12, Mmp14, Spp1, Thbs1, Fn1*) and monocyte-to-macrophage differentiation and maintenance (*Mafb, Bhlhe41, Csf1r, Csf1*). Similarly, genes upregulated in tissue-resident compared with monocyte-derived alveolar macrophages were similar in young and old animals(Figure 6B, Cluster 2) and included canonical genes expressed in mature alveolar macrophages (*Siglecf, Axl, Sirpa, Ear1*) and were enriched for genes related to positive regulation of RNA biosynthesis and histone modification. In agreement with our previous report, genes upregulated in tissue-resident alveolar macrophages with aging (Figure 6B, Cluster 1) were related to antigen processing and presentation and immune system processes, while genes downregulated in tissue-resident alveolar macrophages from old mice (Figure 6B, Cluster 4) were related to positive regulation of cell cycle (39).

### A single dose of ISRIB resets proteostasis in the lung epithelium and modulates stress response in epithelial cells

Previous studies have demonstrated that even a single dose of ISRIB could dramatically ameliorate ISR-driven diseases (28, 29, 40). Accordingly, we treated mice with a single dose of ISRIB or vehicle 7 days after administration of bleomycin and assessed severity of fibrosis 21 day later (Figure 7A). Mice treated with ISRIB reduced histologic evidence of fibrosis as measured by blinded scoring of randomly selected lung sections stained with Masson’s trichrome, improved lung compliance and decreased collagen levels (Figure 7B and C). In addition, we asked whether prophylactic administration of ISRIB is sufficient to ameliorate bleomycin-induced pulmonary fibrosis. We treated mice with a single dose of ISRIB or vehicle immediately prior to bleomycin instillation. Mice treated with ISRIB exhibited less severe fibrosis (Figure 7B and C). Moreover, mice treated with ISRIB lost significantly less weight than vehicle-treated mice (Figure 7D).

**Figure 7.**
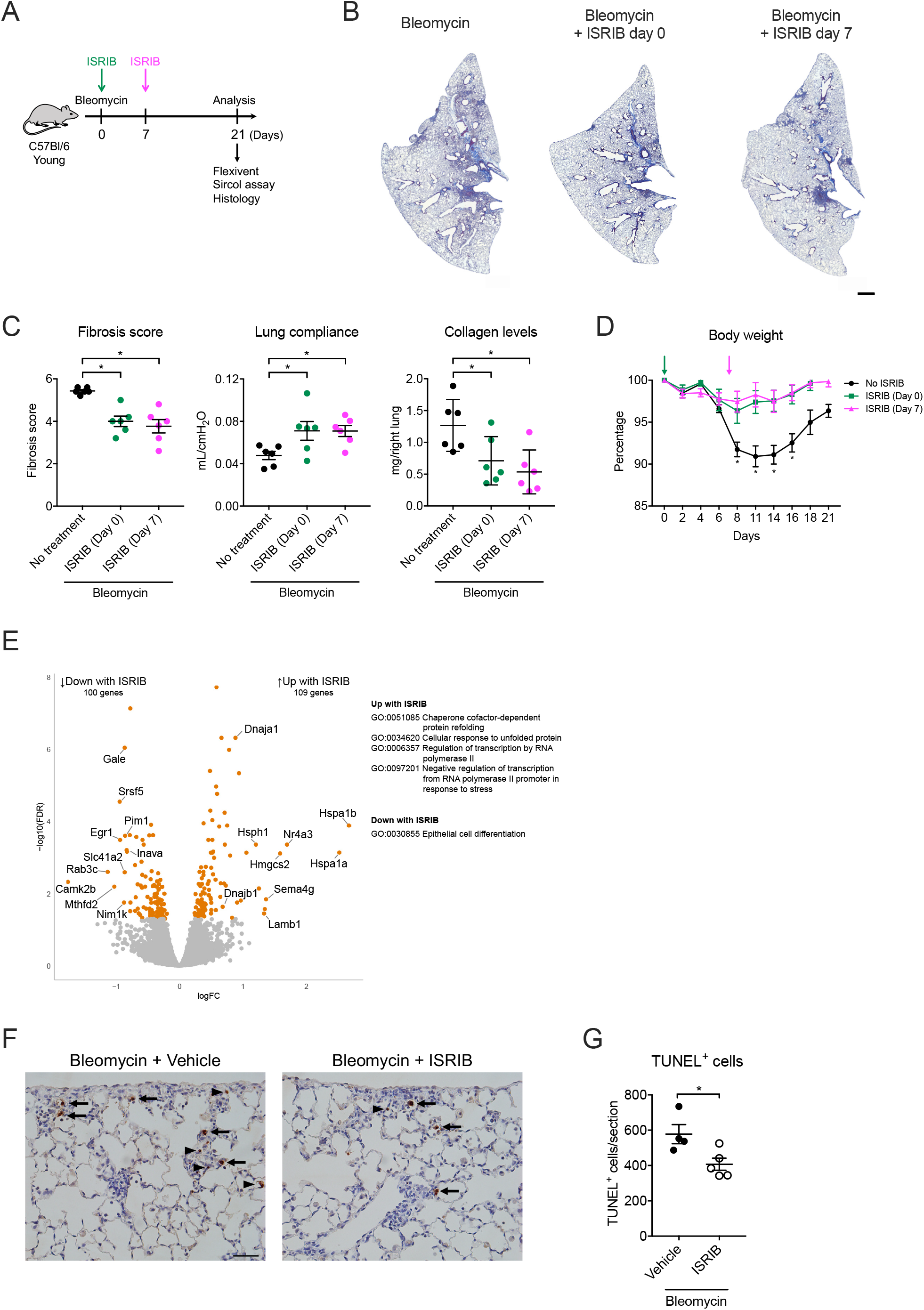
ISRIB ameliorates fibrosis by modulating stress response in epithelial cells. (**A**) Schematic of the experiment outline. Young (3 months) mice were administered 0.025 unit of bleomycin and treated with 25 mg/kg of ISRIB according to the time line. (**B**) Representative histologic images in bleomycin model (Masson’s trichrome, Scale bar = 1 mm). (**C**) Quantification of fibrosis score using Ashcroft score, lung compliance using Flexivent, and soluble collagen in lung homogenates in bleomycin model. (**D**) Body weight in bleomycin model with single dose of ISRIB treatment. Arrow indicates the point of ISRIB treatment. (**E**) Volcano plot showing number of differentially expressed genes in alveolar epithelial cells in naïve young mice 24 hours after vehicle or ISRIB treatment (FDR q-value < 0.05). Characteristic GO processes are shown on the right (p-value < 0.05). (**F**) Representative figures of lung tissue with TUNEL staining from bleomycin-treated young mice at day 7 that were treated with single dose of ISRIB or vehicle 24 hours before. Arrows indicate TUNEL-positive epithelial cells and triangle arrows indicate TUNEL-positive immune cells. Scale bar = 100 µm. (**G**) The number of TUNEL-positive cells per section. Data are shown as mean ± SEM, 4-5 mice per group. One-way ANOVA with Tukey-Kramer test for multiple comparison, non-parametric analysis with Mann-Whitney test, or Kruskal-Wallis test with Dunn’s multiple comparisons test for comparison of each body weight to baseline (day 0). *p < 0.05.

These data suggest that ISRIB ameliorates pulmonary fibrosis by modulating signals upstream of monocyte-derived alveolar macrophage recruitment. As intratracheal bleomycin and asbestos primarily impact the lung epithelium, we treated mice with a single dose of ISRIB or vehicle and 24 hours later performed transcriptomic profiling on flow-sorted alveolar epithelial type II cells: 100 genes were downregulated and 109 genes were upregulated (FDR q < 0.05). The upregulated genes included genes critical to protein folding (*Hspa1a, Hspa1b, Hsph1, Bag3, Dnajb1* and others), while downregulated genes were enriched for biological processes related to cell death and cell differentiation (Figure 7E, Supplemental Table S6, S7). These data suggested that a single dose of ISRIB was sufficient to induce preemptive upregulation of chaperones in epithelial cells, rendering them more resistant to subsequent stress and apoptosis. We and others have shown that genetic strategies that prevent apoptosis reduce the severity of pulmonary fibrosis in response to bleomycin or forced overexpression of TGF-β (41, 42). We administered ISRIB 6 days after the intratracheal instillation of bleomycin and quantified the number of apoptotic cells 24 hours later using TUNEL assay. Treatment with ISRIB significantly reduced the number of TUNEL-positive cells (Figure 7F and G).

## DISCUSSION

Our data add to an emerging literature elucidating the multicellular nature of pulmonary fibrosis. We found that a small molecule inhibitor of the integrated stress response – ISRIB – resets proteostasis in the alveolar epithelium thereby reducing the recruitment of pathogenic monocyte-derived alveolar macrophages. We and others have shown that after recruitment these monocyte-derived alveolar macrophages form stable, perturbation-resistant circuits that drive fibroblast proliferation and matrix production (16–18). Over time these fibroblasts may gain the ability to maintain proliferation and matrix production independent of macrophages, leading to localized areas of unchecked fibrosis (43, 44).

We observed that old mice developed more severe pulmonary fibrosis in response to both bleomycin and asbestos. We used a genetic lineage tracing system that allowed us to measure the dynamics of monocyte-derived alveolar macrophage recruitment over the course of resolving (bleomycin)- and non-resolving (asbestos)-induced fibrosis in young and aged mice. In young mice exposed to bleomycin, monocyte-derived alveolar macrophages were recruited as a single wave of cells confined to the first 14 days after injury. In contrast, more monocyte-derived alveolar macrophages were recruited to the lungs of old mice, and recruitment was ongoing even 21 days after the injury. Consistently, the recruitment of monocyte-derived alveolar macrophages was enhanced at all time points in old compared with young mice after asbestos exposure. Transcriptomic profiling suggested minimal differences in profibrotic gene expression of monocyte-derived alveolar macrophages in old compared with young mice and mice treated with ISRIB. Instead, an apparent increase in fibrotic gene expression in monocyte-derived alveolar macrophages from old mice after bleomycin was attributable to ongoing recruitment of immature cells.

Combined, these lineage tracing and transcriptomic studies suggest that age-related changes in the number and transcriptomic identity of monocyte-derived alveolar macrophages during fibrosis are not cell-autonomous. Instead, these changes represent responses to the aging lung microenvironment. This is in agreement with our previous finding in normally aged mice that orthotopic heterochronic adoptive transfer of tissue-resident alveolar macrophages from young into old mice results in a change of transcriptomic identity that reflects the age of the recipient (39). Because ISRIB reduced the recruitment of monocyte-derived alveolar macrophages to the lungs of old and young mice without affecting their transcriptomic identity, these findings suggest that ISRIB targets the lung epithelium to reduce fibrosis in these models. Indeed, even a single dose of ISRIB given to naïve mice was sufficient to induce the expression of *Hsp1a* and *Hsp1b*, key members of HSP70 chaperone family, and their co-chaperone with a known anti-apoptotic function – *Bag3* (45). Consistently, the administration of ISRIB reduced the levels of apoptosis during bleomycin-induced lung injury. Whether ISRIB reduced the recruitment of monocyte-derived alveolar macrophages by reducing epithelial susceptibility to bleomycin- or asbestos-induced injury, promoting epithelial repair or enhancing the age-related loss of homeostatic signals necessary for alveolar macrophage maturation, is not known.

ISRIB is a small molecule developed by Carmela Sidrauski and colleagues in Peter Walter’s laboratory that targets the integrated stress response (ISR) (29). The ISR is activated in response to environmental or genetic stressors that threaten the proteome including ER stress (via PERK), viral infection (via PKR), amino acid deprivation (via GCN2) or heme deprivation (via HRI) (12). These kinases all phosphorylate eIF2α, a component of eukaryotic initiation factor 2 (eIF2), which is required to initiate translation on AUG start codons as a part of eIF2-GTP-Met-tRNA ternary complex. During translation initiation GTP is hydrolyzed to GDP, and a dedicated guanine nucleotide exchange factor for eIF2 (eIF2B) is needed to catalyze GDP-to-GTP exchange and reactivate eIF2 (44). Phosphorylation of eIF2α causes it to bind tightly to eIF2B precluding binding of the unphosphorylated protein and preventing binding to GTP (46). This inhibits bulk translation but enhances translation of proteins with small upstream open reading frames. Important among these is ATF4, a transcription factor that drives the expression of proteins that protect the proteome including chaperones and metabolic enzymes among others (47). ISRIB enhances guanine exchange factor activity of eIF2B in the presence of phosphorylated eIF2α, relieving both translational inhibition and gene expression induced by ATF4 (29). Like many stress responses, the activation of ATF4 exhibits hormesis; short-term activation is cytoprotective while longer-term or very high level activation is detrimental to the cell (48). Indeed, ATF4 induces the transcription of CHOP, which induces apoptosis upon reaching a threshold level (49). Therefore, ISRIB might function to reduce ISR and promote an adaptive response and survival short-term, while being detrimental during the chronic stress (50).

Our finding that ISRIB attenuates bleomycin- and asbestos-induced fibrosis in young and old mice adds to a growing body of literature showing dramatic effects of ISRIB in age-related disease. A single dose of ISRIB has been shown to improve cognitive function in normal mice and in mice after traumatic brain injury (28, 29, 51), prevent neurodegeneration in murine prion disease (52), improve glucose tolerance and diabetes induced liver injury in rats (53), prevent postnatal hearing loss in a genetic mouse model (54), reduce neuropathic pain in a murine model of diabetic neuropathy (55), alleviate dwarfism in a murine model of chondrodysplasia (56), slow the growth of metastatic prostate cancer (57), improve survival of lymphoblast cell lines from patients with vanishing white matter disease (58), and alleviate the social deficit and elevated anxiety-like behavior in a murine model of these neuropsychiatric disorders (59). Interestingly, while ISRIB improved memory in normal animals, it did not do so in two murine models of Alzheimer’s disease (60, 61).

The finding that ISRIB targets the epithelium to prevent lung fibrosis is consistent with genetic evidence implicating proteostatic stress in the initiation of pulmonary fibrosis (62). For example, a mutation in the *MUC5B* promoter associated with fibrosis results in enhanced expression of an abundant mucin secreted by lung epithelial cells (25). A mutation in *SFTPC*, the expression of which is restricted to alveolar type II cells, encodes a misfolded protein and expression of this mutant in alveolar type II cells from mice is sufficient to induce spontaneous pulmonary fibrosis (20, 21, 24). Loss of function in genes associated with the Hermansky– Pudlak syndrome disrupt intracellular protein transport and investigators have localized the effect of these mutants to the lung epithelium in animal models (22, 23, 63). While genes involved in the maintenance of telomere length are ubiquitously expressed, loss of function mutations in these genes targeted to alveolar type II cells induce spontaneous or more severe fibrosis in animal models (64, 65). Consistent with these findings, genetic and pharmacologic strategies to inhibit ER stress have been reported to reduce bleomycin-induced fibrosis (34, 36, 66).

Our study has several limitations. First, while we showed transcriptomic changes in the lung epithelium, reduced apoptosis and reduced numbers of recruited monocyte-derived alveolar macrophages in response to administration of ISRIB during the development of the fibrosis, ISRIB might also have effects on lung repair, during the resolution phase. Second, our finding that age-related increase of the number of lung monocytes was ameliorated by ISRIB treatment suggest it might have effects in the bone marrow or other organs that might secondarily protect the epithelium. Finally, ISRIB both relieves translational inhibition and inhibits the expression of ATF4-mediated gene expression. Whether one or both of these is required for the reduction in monocyte-derived macrophage recruitment and prevention of fibrosis will require additional study.

In conclusion, we show that ISRIB, a small molecule inhibitor of the ISR, can attenuate resolving fibrosis induced by bleomycin and non-resolving fibrosis induced by asbestos fibers in both young and old mice. ISRIB resets proteostasis to enhance the stress resistance of the alveolar epithelium, reducing the recruitment of monocyte-derived alveolar macrophages. ISRIB shows promise as part of a combined strategy targeting epithelial proteostasis, pathogenic monocyte-derived alveolar macrophages and proliferative fibroblasts to treat pulmonary fibrosis.

## MATERIALS AND METHODS

### Key Resources Table

| REAGENT or RESOURCE | SOURCE | IDENTIFIER |
| --- | --- | --- |
| **Antibodies** |  |  |
| Rat anti-mouse I-A/I-E (MHC II), BUV395 | BD Biosciences | Cat# 743876, RRID:AB_2741827 |
| Rat anti-mouse CD24, BUV496 | BD Biosciences | Cat#564664, RRID:AB_2716853 |
| Rat anti-mouse CD11b, BUV737 | BD Biosciences | Cat#564443, RRID:AB_2738811 |
| Rat anti-mouse Ly-6C, eFluor 450 | eBioscience | Cat#48-5932-82, RRID:AB_10805519 |
| Mouse anti-mouse CD64, PE | BioLegend | Cat#139303, RRID:AB_10613467 |
| Mouse anti-mouse NK-1.1, PE-CF594 | BD Biosciences | Cat#562864, RRID:AB_2737850 |
| Hamster anti-mouse CD11c, PE-Cy7 | BD Biosciences | Cat#558079, RRID:AB_647251 |
| Rat anti-mouse Siglec F, Alexa Fluor 647 | BD Biosciences | Cat#562680, RRID:AB_2687570 |
| Rat anti-mouse CD45, APC-Cy7 | BD Biosciences | Cat#557659, RRID:AB_396774 |
| Rat anti-mouse Ly-6G, Alexa Fluor 700 | BD Biosciences | Cat#561236, RRID:AB_10611860 |
| Armenian hamster anti-mouse CD3e, FITC | eBioscience | Cat#11-0031-82, RRID:AB_464882 |
| Rat anti-mouse CD25, PE | eBioscience | Cat#12-0251-82, RRID:AB_465607 |
| Rat anti-mouse CD8a, PE | BioLegend | Cat#100762, RRID:AB_2564027 |
| Rat anti-mouse CD62L, APC-eFluor 780 | eBioscience | Cat#47-0621-82, RRID:AB_1603256 |
| Rat anti-mouse CD19, Alexa Fluor 700 | eBioscience | Cat#56-0193-82, RRID:AB_837083 |
| Rat anti-mouse CD4, BUV395 | BD Biosciences | Cat#563790, RRID:AB_2738426 |
| Rat anti-mouse CD44, BUV737 | BD Biosciences | Cat#564392, RRID:AB_2738785 |
| **Chemicals, Peptides, and Recombinant Proteins** |  |  |
| Bleomycin | Fresenius Kabi | Cat#0703-3154-01 |
| Asbestos (UICC crocidolite) | Provided by US Environmental Protection Agency |  |
| ISRIB | AdooQ<br>BioScience | Cat#A14302 |
| **Critical Commercial Assays** |  |  |
| EdU | ThermoFisher<br>Scientific | Cat#C10044 |
| Click-iT Plus EdU Alexa Fluor 647 Flow<br>Cytometry Assay Kit | ThermoFisher<br>Scientific | Cat#C10634 |
| PicoPure RNA Isolation Kit | ThermoFisher<br>Scientific | Cat#KIT0204 |
| AllPrep DNA/RNA Mini Kit | Qiagen | Cat#80204 |
| NEBNext RNA Ultra I kit | New England<br>Biolabs | Cat#E7530L |
| polyA mRNA isolation module | New England<br>Biolabs | Cat#E7490S |
| SMARTer Stranded Total RNA-Seq Kit v2 -<br>Pico Input Mammalian kit | Takara Bio | Cat#634413 |
| **Deposited Data** |  |  |
| **Experimental Models: Organisms/Strains** |  |  |
| Mouse: C57BL/6J (8-12 weeks old) | The Jackson<br>Laboratory | JAX: 000664 |
| Mouse: C57BL/6J (over 18 months old) | National Institute of Aging |  |
| Mouse: *Cx3cr1<sup>ERCre</sup>* | The Jackson Laboratory | JAX: 020940 |
| Mouse: ZsGreen | The Jackson Laboratory | JAX: 007906 |
| Software and Algorithms |  |  |
| ImageJ | Schneider et al., 2012 | https://imagej.nih.gov/ij/ |
| bcl2fastq 2.19.1 |  | https://support.illumina.com/sequencing/sequencing_software/bcl2fastq-conversion-software.html |
| FastQC |  | https://www.bioinformatics.babraham.ac.uk/projects/fastqc/ |
| Trimmomatic 0.36 |  | http://www.usadellab.org/cms/?page=trimmomatic |
| STAR 2.6.0 | Dobin et al., 2013 | https://github.com/alexdobin/STAR |
| htseq-count 0.11.2 | Anders et al., 2014 | https://htseq.readthedocs.io/en/latest/ |
| edgeR 3.28.0 | Robinson, 2010 |  |
| Other |  |  |

### Methods Details

#### Mice

All experimental protocols were approved by the Institutional Animal Care and Use Committee at Northwestern University (Chicago, IL, USA). All strains including wild-type mice are bred and housed at a barrier- and specific pathogen–free facility at the Center for Comparative Medicine at Northwestern University. Eight to twelve weeks old mice were used as young mice and over eighteen months old mice were used as old mice for all experiments. The C57BL/6J mice were obtained from Jackson laboratories (Jax stock 000664). Old mice were obtained from National Institute of Aging. The *Cx3cr1*^*ERCre*^ mice (67) and *ZsGreen* (68) mice were obtained from Jackson laboratories (Jax stocks 020940 and 007906, correspondingly).

#### Bleomycin- and asbestos-induced lung fibrosis and drug administration

Mice were anesthetized and intubated followed by intratracheally instillation of bleomycin (0.025 unit in 50 µl PBS) or crocidolite asbestos fibers (100 µg in 50 µl PBS) to induce lung-fibrosis, as previously described (16, 18). Lungs were harvested at indicated time points for flow cytometry, RNA sequencing, and histopathology. ISRIB (AdooQ BioScience, A14302) was first reconstituted in DMSO to the concentration 4 mg/ml, and then further diluted using phosphate buffered saline to the final concentration 0.3125 mg/ml (52). ISRIB was administered via intraperitoneal injection, either via single or daily injections, as indicated in the text and figure legends, at the dose 2.5 mg/kg in a final volume of 200 ul. Control animals were treated using vehicle.

#### Tissue preparation, flow cytometry and sorting

Tissue preparation for flow cytometry analysis and cell sorting was performed as previously described (16, 18), with modifications. Briefly, mice were euthanized and their lungs were perfused through the right ventricle with 10 ml of HBSS. The lungs were removed and infiltrated with 2 mg/ml collagenase D (Roche, Indianapolis, Indiana) and 0.2 mg/ml DNase I (Roche, Indianapolis, Indiana) dissolved in HBSS with Ca^2+^ and Mg^2+^, using syringe with 30G needle. Lungs were chopped with scissors, tissue was transferred into C-tubes (Miltenyi Biotech, Auburn, California, 130-096-334), and processed in a GentleMACS dissociator (Miltenyi Biotech, Auburn, California) using program *m_lung_01*, followed by incubation for 30 minutes at 37°C with gentle agitation, followed by program *m_lung_02*. The resulting single-cell suspension was filtered through a 40 µm nylon cell strainer. The cells were incubated with anti-mouse CD45 microbeads (Miltenyi Biotech, Auburn, California,130-052-301) and CD45^+^ cells were collected using the MultiMACS Cell24 Separator (Miltenyi Biotech, Auburn, California), using *possel* protocol. Automated cell counting was performed using Nexcelom K2 Cellometer with AO/PI reagent. Cells were stained with fixable viability dye eFluor 506 (eBioscience, Waltham, Massachusetts), incubated with FcBlock (BD Biosciences, San Jose, California), and stained with a mixture of fluorochrome-conjugated antibodies, as listed above. Click-it reaction for Alexa Fluor 647 labelling were performed using the Click-iT Plus EdU Alexa Fluor 647 Flow Cytometry Assay Kit (Thermo Fisher Scientific, C10634). Single color controls were prepared using BD CompBeads (BD Biosciences, San Jose, California) and Arc beads (Invitrogen, Waltham, Massachusetts). Flow cytometry and cell sorting were performed at the Northwestern University Robert H. Lurie Comprehensive Cancer Center Flow Cytometry Core facility (Chicago, Illinois). Data were acquired on a custom BD FACSymphony instrument using BD FACSDiva software (BD Biosciences, San Jose, California). Compensation and analysis were performed using FlowJo software (TreeStar). Each cell population was identified using sequential gating strategy (Figure S2). The percentage of cells in the live/singlets gate was multiplied by the number of live cells using Cellometer K2 Image cytometer to obtain cell counts. Cell sorting was performed using BD FACSAria III instrument using 100 um nozzle and 40 psi pressure.

#### Bulk RNA-seq

RNA was isolated using Arcturus PicoPure RNA Isolation Kit (Thermo Fisher Scientific, catalog number KIT0204) for experiments in Figures 6A, 7C or AllPrep DNA/RNA Mini Kit (Qiagen, catalog number 80204) for experiment in Figure 6B. RNA quality was assessed using TapeStation 4200 (Agilent, Santa Clara, California), only samples with RNA integrity number over 7 were used for library preparation. Library construction for RNA-seq was performed using NEBNext RNA Ultra I kit (New England Biolabs, catalog number E7530L) with polyA mRNA isolation module (New England Biolabs, catalog number E7490S) from 30ng (Figure 6B), 3ng (Figure 6A) or 50ng (Figure 7C) of RNA. Libraries were assessed for quality (TapeStation 4200, Agilent, Santa Clara, California) and then sequenced on NextSeq 500 instrument (Illumina, San Diego, California). FASTQ files were generated using bcl2fast (version 2.19.1), followed by quality control using FastQC, trimming using Trimmomatic (version 0.36) and mapping to mm10 version of the mouse genome with STAR (version 2.6.0). Counts were generated using htseq-count (HTSeq framework, version 0.11.2). Differential gene expression was performed using edgeR (version 3.28.0). Raw data are available from GEO: GSE145295, GSE145590, GSE145771. Detailed R code used for analysis is available at https://github.com/NUPulmonary/Watanabe2020.

#### Histopathology and immunohistochemistry

After euthanasia and perfusion, trachea was cannulated with Luer syringe stub blunt needle and mouse lungs were inflated with 4% paraformaldehyde at 15 cm H_2_O column pressure. Left lung was fixed in 4% paraformaldehyde for 24 hours, dehydrated and embedded in paraffin. For histopathology 4 µm thick sections were prepared. Hematoxylin and eosin staining and Masson’s trichrome staining were performed for the analysis of fibrosis scoring. TUNEL assay was performed for the analysis of apoptotic cells. Immunohistochemistry was performed at Northwestern University Mouse Histology and Phenotyping Laboratory Core facility (Chicago, Illinois).

#### Fibrosis scores and lung collagen determination

Fibrosis scores were performed from Masson’s trichrome– stained specimens in a blinded manner, in accordance with modified Ashcroft score for bleomycin (69) and the code set by pathology standard for asbestosis (70). Collagen levels were determined using Sircol assay, as described previously (16, 18). For collagen levels using second harmonic generation, the deparaffinized mouse lung tissue were imaged on Nikon A1R-MP Multiphoton Microscope at Northwestern University Nikon Cell Imaging Facility (Chicago, Illinois).

#### Statistical analysis

Statistical tests and tools for each analysis are explicitly described with the results or detailed in figure legends.

---

**Figure S1.**
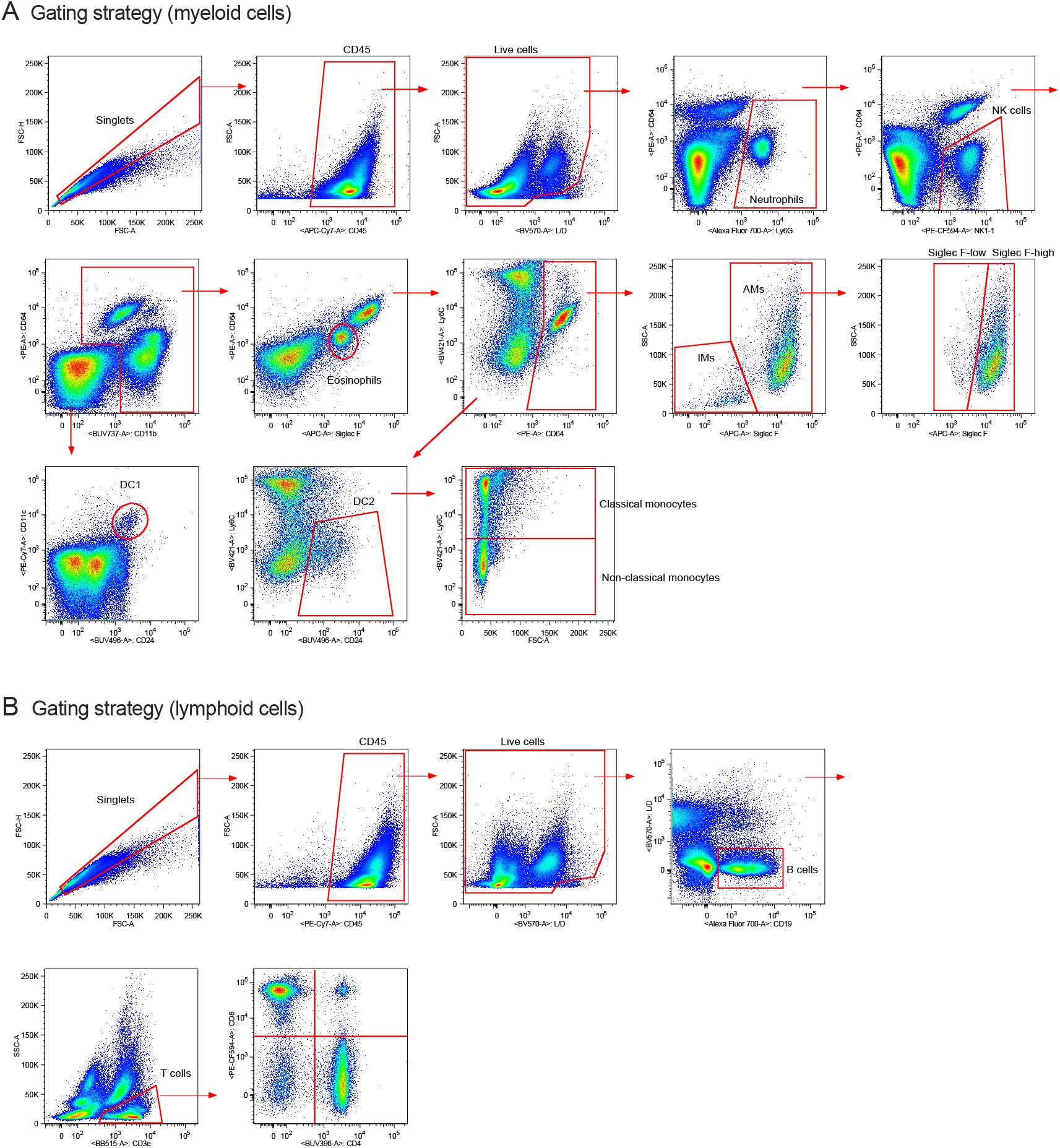
Flow cytometry gating strategy for lung myeloid and lymphoid cell populations. Gating strategy to enumerate and flow cytometry sort (**A**) myeloid cell populations and (**B**) lymphoid cell populations. Representative data from naïve young mouse lung are shown.

**Figure S2.**
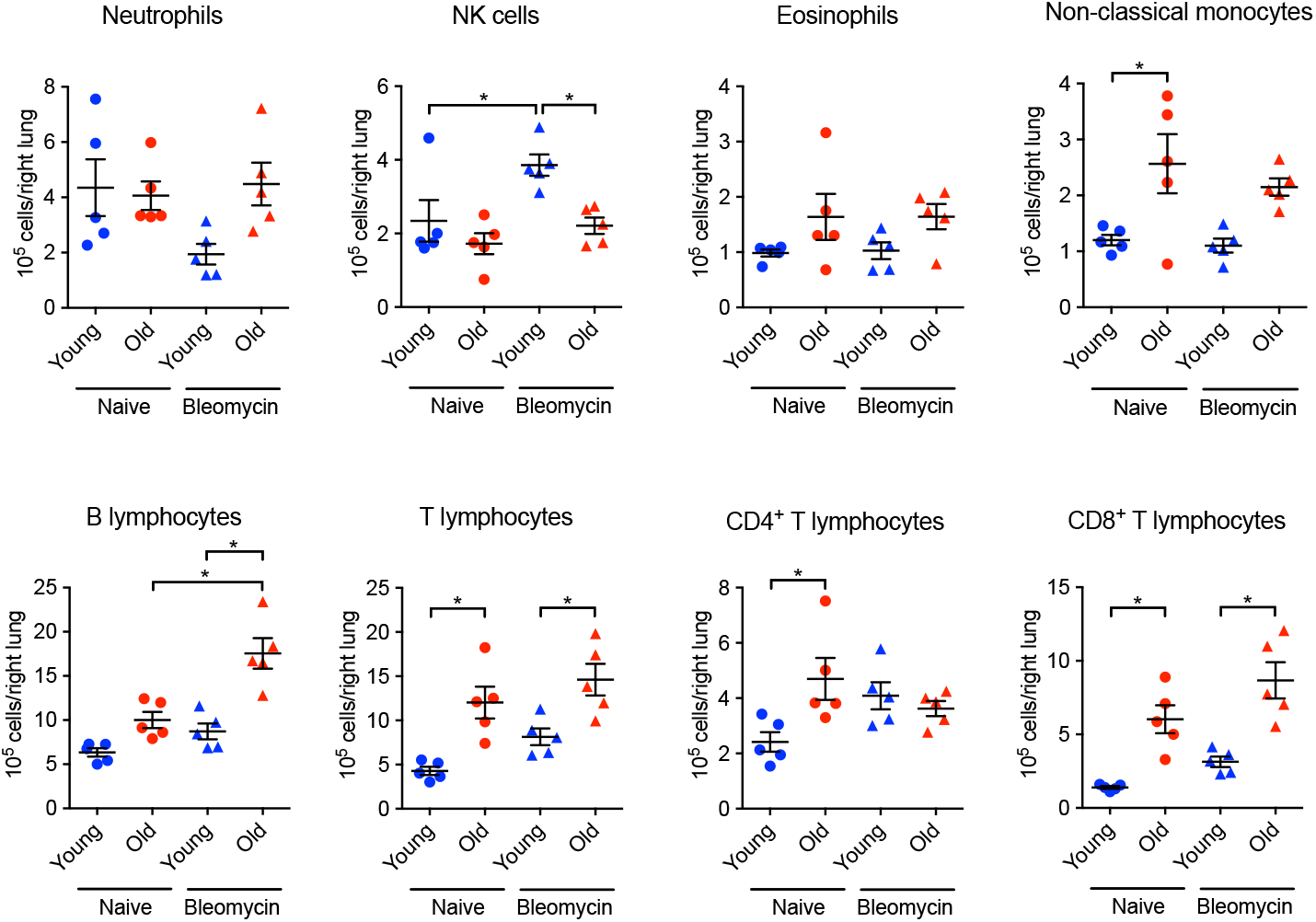
Aging is associated with the changes in the number of lymphoid and myeloid cell populations in naïve mice and during lung fibrosis. Myeloid populations including neutrophils, NK cells, eosinophils and non-classical monocytes and lymphoid populations including B cells, T cells, CD4^+^ T cells and CD8^+^ T cells were enumerated using flow cytometry. Data presented as mean ± SEM, 5–8 mice per group, one-way ANOVA with Tukey-Kramer test for multiple comparisons. *, p< 0.05.

**Figure S3.**
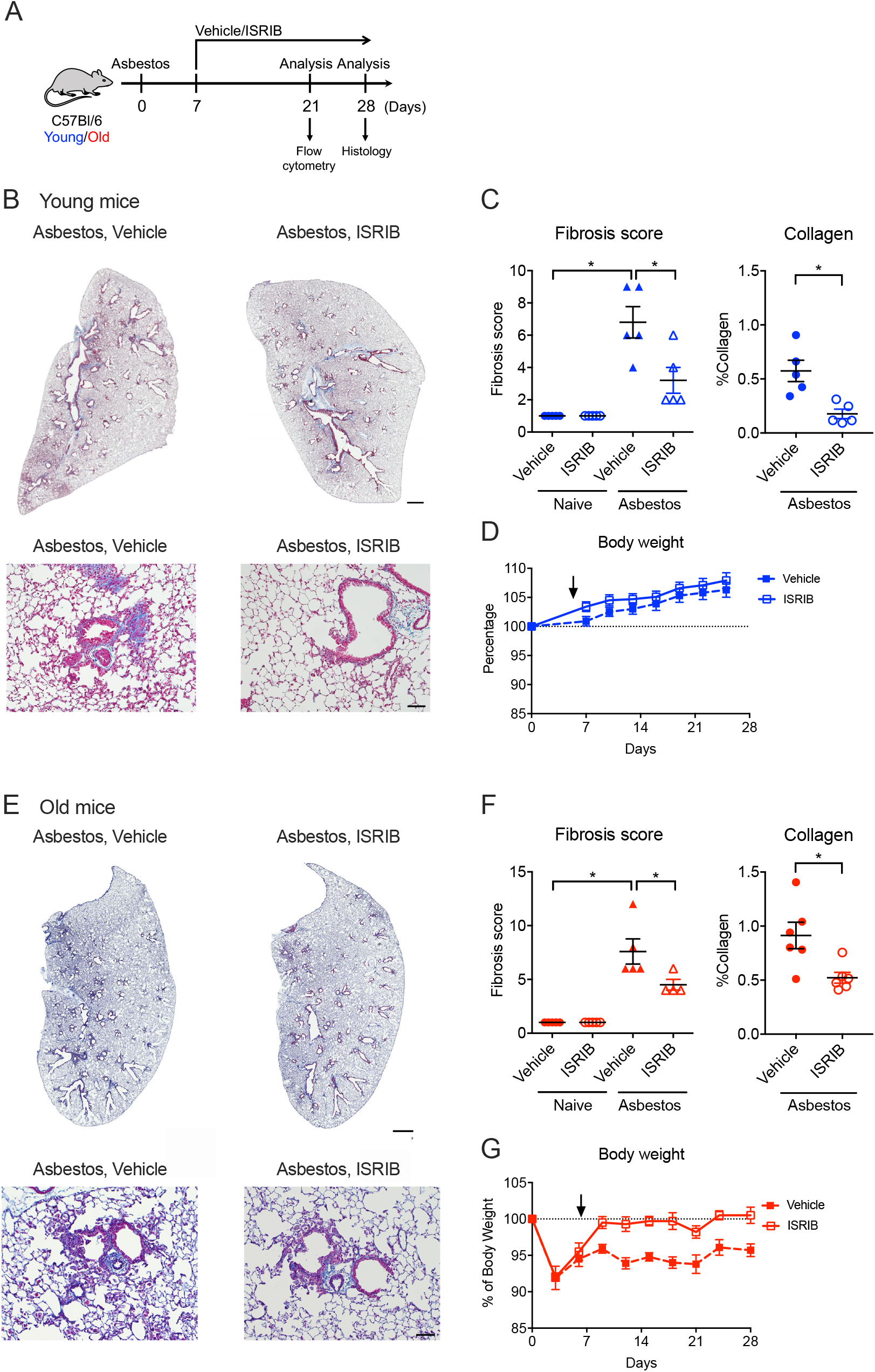
Therapy with ISRIB attenuates asbestos induced pulmonary fibrosis in young and old mice. (**A**) Schematic of the experiment design. Young (3 months) and old (over 18 months) mice were administered crocidolite asbestos (100 μg/50 µl, intratracheally) and treated with 25 mg/kg of ISRIB or vehicle (i.p., every day) beginning at Day 7 when fibrosis has begun and harvested at the indicated time points. (**B, E**) Representative images of lung tissue from naïve mice or asbestos-treated mice with or without ISRIB on day 28 and fibrosis score. Masson’s trichrome staining. Scale bar = 1 mm. Representative images in high magnification (400×) are also shown. Scale bar = 100 μm. (**C, F**) Fibrosis score and the percentage of collagen levels using secondary harmonic generation from naïve mice or asbestos-treated mice with or without ISRIB. (**D, G**) The graph shows the body weight. Arrow indicates the starting point of ISRIB treatment. Data are shown as mean ± SEM, 5-7 mice per group. One-way ANOVA with Tukey-Kramer test for multiple comparison, non-parametric analysis with Mann-Whitney test, or Kruskal-Wallis test with Dunn’s multiple comparisons test for comparison of each body weight to baseline (day 0). *p < 0.05. Representative data from two independent experiments.

**Figure S4.**
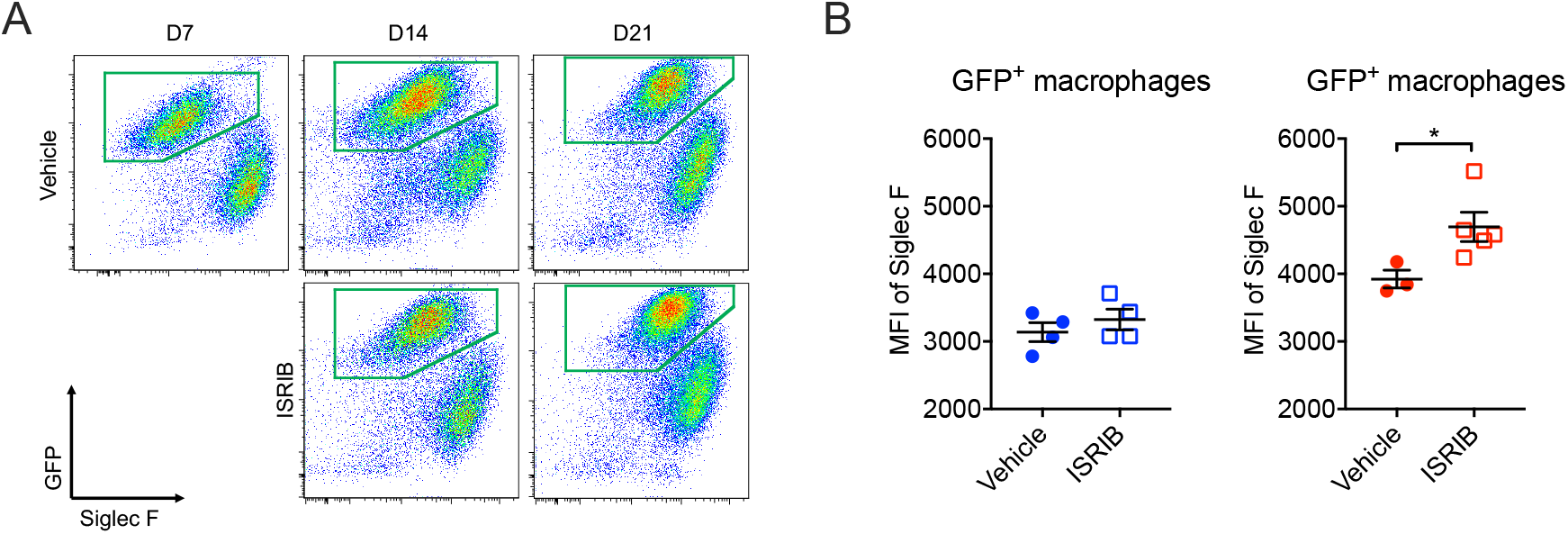
ISRIB promotes the maturation of monocyte-derived alveolar macrophages in old mice during pulmonary fibrosis. (**A**) Representative flow data gated on CD45^+^CD64^+^ macrophages. (**B**) Median fluorescence intensity (MFI) of Siglec F levels on GFP^+^ macrophages in young and old mice 21 days after bleomycin model with and without ISRIB treatment. Data are shown as mean ± SEM, 3-5 mice per group. Non-parametric analysis with Mann-Whitney test. *p < 0.05.

**Figure S5.**
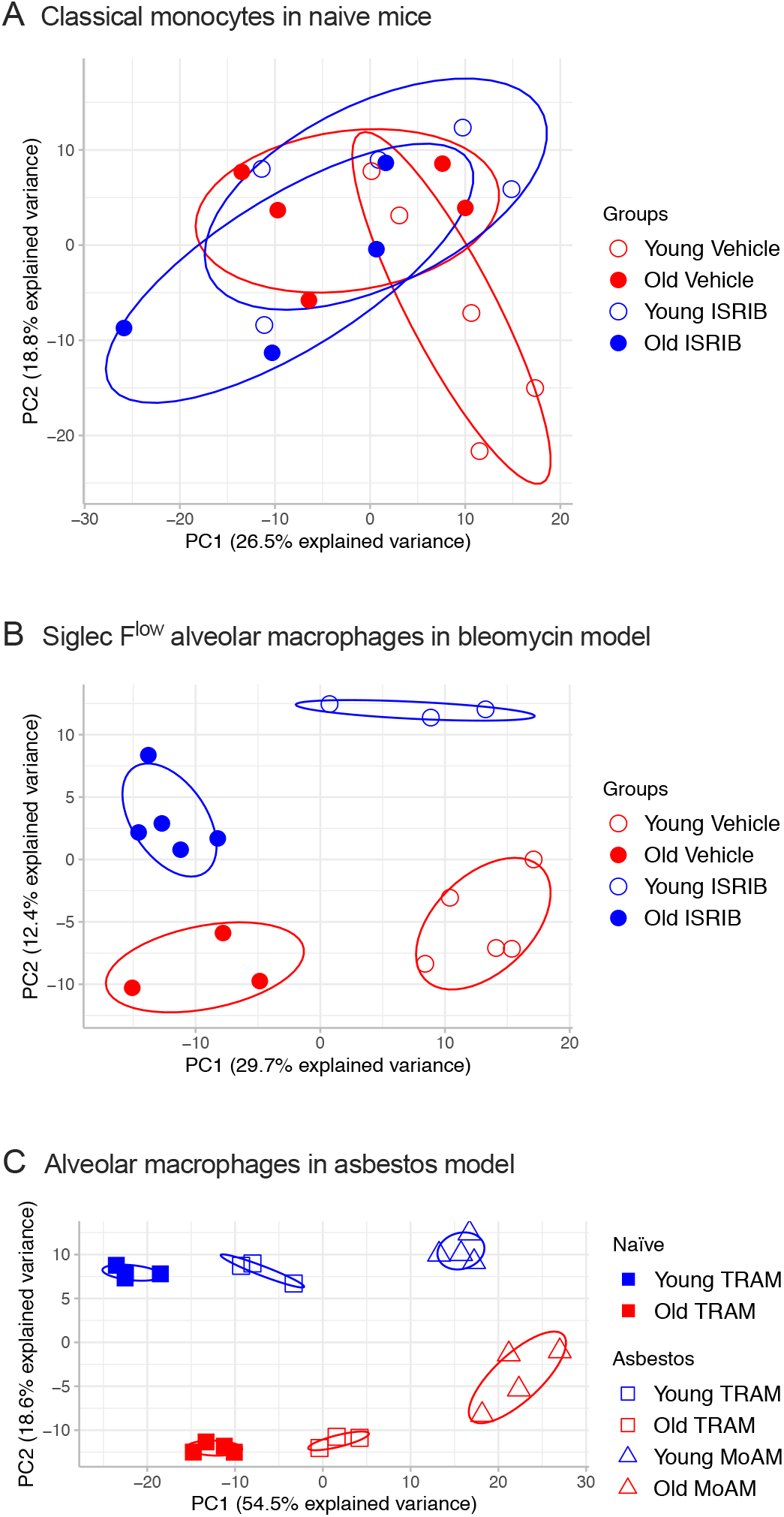
Principal component analysis. (**A**) Principal component analysis of classical monocytes in young and old naïve mice treated with ISRIB or vehicle did not reveal age- or treatment-correlated sources of variation. (**B**) Principal component analysis of monocyte-derived alveolar macrophages in young and old mice bleomycin model with and without ISRIB treatment. PC1 is associated with aging and PC2 is associated with ISRIB treatment. (**C**) Principal component analysis of tissue-resident alveolar macrophages (TRAMs) and monocyte-derived alveolar macrophages (MoAMs) in naïve lung and asbestos-induced lung fibrosis. PC1 is associated with pulmonary fibrosis and PC2 is associated with aging.

## LIST OF SUPPLEMENTAL TABLES

**Supplemental table S1:** Number of differentially-expressed genes between young/old ISRIB/vehicle samples of sorted monocytes. Related to Figure S6A.

**Supplemental table S2:** List of differentially-expressed genes, their clusters and statistics from Figure 6A.

**Supplemental table S3:** List of GO biological processes for each cluster in Figure 6A.

**Supplemental table S4:** List of differentially-expressed genes, their clusters and statistics from Figure 6B.

**Supplemental table S5:** List of GO biological processes for each cluster in Figure 6B.

**Supplemental table S6:** List of differentially-expressed genes and their statistics from Figure 7C.

**Supplemental table S7:** List of GO biological processes for up- and down-regulated genes in Figure 7C.

## Acknowledgments

This work used services from Northwestern University Flow Cytometry Facility, Center For Advanced Microscopy, and Pathology Core Facility, which are supported by NCI Cancer Center Support Grant P30 CA060553 awarded to the Robert H. Lurie Comprehensive Cancer Center. Multiphoton microscopy was performed on a Nikon A1R multiphoton microscope, acquired through the support of NIH 1S10OD010398-01. This research was supported in part through the computational resources and staff contributions provided by the Genomics Computing Cluster (Genomic Nodes on Quest) which is jointly supported by the Feinberg School of Medicine, the Center for Genetic Medicine, and Feinberg’s Department of Biochemistry and Molecular Genetics, the Office of the Provost, the Office for Research, and Northwestern Information Technology.

## Author contributions

Satoshi Watanabe: designed the study, performed experiments, analyzed results, wrote manuscript. Nikolay S. Markov: performed bioinformatics analysis, wrote manuscript. Ziyan Lu, Raul Piseaux Aillon, Saul Soberanes, Constance E. Runyan, Ziyou Ren, Rogan A. Grant, Mariana Maciel, Hiam Abdala-Valencia, Yuliya Politanska, Kiwon Nam, Lango Sichizya, Hermon G. Kihshen, Nikita Joshi, Alexandra C. McQuattie-Pimentel, Paul A. Reyfman: performed experiments, analyzed results. Richard I. Morimoto: designed and supervised the study, performed analysis, wrote manuscript. GR Scott Budinger and Alexander V. Misharin: designed and supervised the study, performed analysis, wrote manuscript, provided funding for the project. All authors discussed the results and commented on the manuscript.

